# Structural determinants and genetic modifications enhance BMP2 stability and extracellular secretion

**DOI:** 10.1101/373654

**Authors:** Vinayak Khattar, Joo Hyoung Lee, Hong Wang, Soniya Bastola, Selvarangan Ponnazhagan

## Abstract

The short half-life and use of recombinant bone morphogentic protein (BMP)-2 in large doses poses major limitations in the clinic. Events regulating post-translational processing and degradation of BMP2 *in situ*, linked to its secretion, have not been understood. Towards identifying mechanisms regulating intracellular BMP2 stability, we first discovered that inhibiting proteasomal degradation enhances both intracellular BMP2 level and its extracellular secretion. Next, we identified BMP2 degradation occurs through an ubiquitin-mediated mechanism. Since ubiquitination precedes proteasomal turnover and mainly occurs on lysine residues of nascent proteins, we systematically mutated individual lysine residues within BMP2 and tested them for enhanced stability. Results revealed that substitutions on four lysine residues within the pro-BMP2 region and three in the mature region increased both BMP2 turnover and extracellular secretion. Structural modeling revealed key lysine residues involved in proteasomal degradation occupy a lysine cluster near proprotein convertase cleavage site. Interestingly, mutations within these residues did not affect biological activity of BMP2. These data suggest preventing intracellular proteasomal loss of BMP2 through genetic modifications can overcome limitations related to its short half-life.

## INTRODUCTION

Demographic data reveals that steadily rising age of the population in many developed countries is being accompanied by increasing number of skeletal defects every year including an estimated 6.8 million fractures in U.S. alone each year [1, 2]. Autologous and allogeneic bone grafts are two commonly employed strategies to treat bone degeneration [3]. However, the lack of availability of high-quality bone material limits the use of autologous bone repair whereas allogenic bone grafts are complicated by immunosuppression [3]. Discovery of bone morphogenetic protein (BMP)-2 was a significant step to treating bone fractures as recombinant BMP2 is currently being used widely for managing different types of skeletal injuries [4]. BMP2 is a secreted ligand of TGFβ superfamily that plays an indispensable role in osteogenesis [5]. BMP2 is synthesized as an inactive 396 amino acid long pre-pro-protein, which is then proteolytically processed to generate a disulfide-linked homodimer, composed of two monomers of 114 amino acid each [5]. This mature ligand then binds to BMP receptors leading to activation of Smad-dependent signaling pathways and activation of critical genes that trigger bone formation [5]. Despite the potential of BMP2 in clinical applications, its short half-life presents a major problem, necessitating repeated use in large doses of the purified protein at enormous cost and accompanying risks of heterotopic ossification, hematomas, inflammatory reactions and secondary infections within the fracture sites [4, 6-10]. *In vivo* studies have shown a half-life of BMP2 in circulation is very short, averaging 7-16 min systemically [11]. To overcome issues related to its sub-optimal delivery, retention scaffolds are coated with supra-physiological amounts of the protein [7, 8, 12].

Significant research are being undertaken to minimize the rapid loss of BMP2 during therapeutic application, including use of efficient retention scaffolds, gene-delivery methods, crosslinking approaches, and small molecule adjuvants to enhance BMP2 retention, secretion, and synthesis [7, 13, 14]. Some of these studies have also focused on improving BMP2 stability in solution, on retention scaffolds and in the circulation [7, 13]. In contrast, almost no effort has been given to identify mechanisms regulating intracellular abundance of BMP2 and how *in situ* degradation pathways influence its extracellular secretion and bioavailability.

Degradation through the ubiquitination-mediated proteasomal pathway is a cellular mechanism for regulating protein abundance, including those involved in osteogenic differentiation [15, 16]. For instance, specific E3 ubiquitin ligases can regulate osteoblast differentiation by orchestrating the abundance of proteins such as Smad, Runx2, and JunB through the ubiquitination-mediated proteasomal degradation pathway [16]. Ubiquitination is regulated through a wide variety of post-translational modifications, including phosphorylation and glycosylation. BMPs are also subjected to different types of post-translational modification. including glycosylation and phosphorylation [17-19]. More importantly, glycosylation of BMP1 and −2 is known to regulate their stability and secretion [17, 18]. BMP1, lacking critical glycosylation sites, is translocated to the proteasome for degradation [18]. High-throughput proteomic studies have suggested that other members of this family, such as TGFβ1, BMP3, BMP4, BMP6, BMP7, and BMP8 are ubiquitinated in different cell types and tissues [20-22].

It remains to be determined if BMP2 turnover is directly regulated by ubiquitination-mediated proteasomal turnover. This becomes particularly significant as proteasomal inhibitors have been reported to stimulate bone formation both *in vitro* and *in vivo,* although underlying mechanism remains unknown [23, 24]. We surmised that understanding mechanisms regulating intracellular turnover of BMP2 and its effects on secretion would be important for developing strategies to maximize bioavailability, especially for cell- and gene-based applications. To this end, we examined the intracellular degradation kinetics of human BMP2 following perturbation of proteasomal pathway and found that pharmacological blockade of this pathway significantly improved intracellular retention of BMP2 and concomitantly enhanced its secretion. Furthermore, we identified that following synthesis, BMP2 is regulated by ubiquitination-mediated turnover. Since ubiquitination primarily occurs on lysine residues, we individually mutated lysine residues on both pro-BMP2 and mature BMP2 and identified seven key lysine residues, six of which form a lysine cluster at the proprotein convertase cleavage site of BMP2, and which when mutated to arginine increased BMP2 intracellularly and led to a significantly enhanced its secretion without affecting the ligand function.

## RESULTS

### Blocking of proteasomal degradation pathway leads to a dose-dependent and time-dependent increase in BMP2 level *in-situ* and leads to enhanced secretion

To identify if proteasomal blockade affects BMP2 regulation, we assessed intracellular and secreted BMP2 levels following treatment of BMP2-transfected 293T cells with two different proteasomal inhibitors: epoximicin and MG-132. Epoximicin is a highly potent, irreversible and one of the most specific proteasome inhibitors that also primarily reacts with the chymotrypsin-like site [25, 26]. MG-132 is a reversible and cell-permeable inhibitor that primarily acts on the chymotrypsin-like site in the β subunit of the proteasome [27]. 293T cells were transfected with HA-tagged BMP2 expression plasmid, pCMV3-HA-BMP2. Following transfection, cells were treated with increasing doses of epoximicin or MG-132 for 6 hours and both intracellular and secreted BMP2 level was assessed by immunoblotting. Results of the study indicated a significant dose-dependent increase in intracellular BMP2 level following treatment with epoximicin and MG-132, compared to control (**P≤0.01, *P≤0.05; **Figure 1A**, and **Supp. Figure 1 A**, upper panel). The increase in BMP2 level *in-situ* following proteasomal blockade was also accompanied by a corresponding increase in BMP2 secretion (**P≤0.01 to *P≤0.05; **Figure 1B**, and **Supp. Figure 1A**, lower panel).

**Figure 1:**
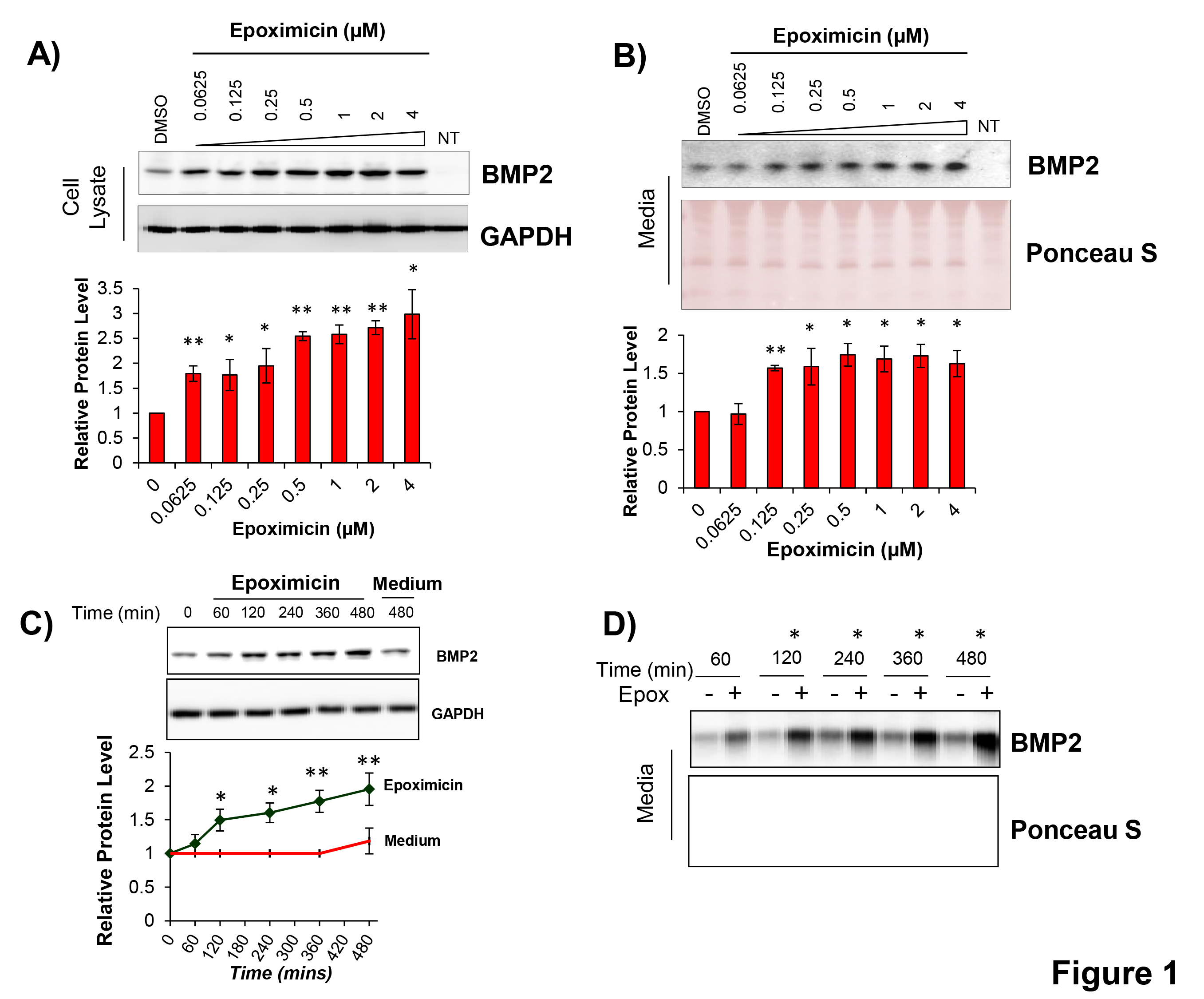
Effect of epoximicin on intracellular BMP2 level and extracellular secretion. 293T cells were transfected with a mammalian expression vector encoding BMP2. The cells were split 24 hrs post-transfection and treated with increasing concentration of epoximicin. The cells were harvested and both cell lysate and media were collected 6 hours post treatment. Levels of intracellular (**A**) and secreted (**B**) BMP2 were determined by immunoblotting. Densitometric quantitation from triplicate experiment for intracellular and secreted BMP2 levels following treatment with epoximicin is shown below each blot (*p<0.05; **p<0.01, compared to media alone). In replicate experiments, following transfection of BMP2 expression vector, cells were treated with 2 μM epoximicin for increasing duration of time (**C**). Cells and media were harvested at regular intervals, and BMP2 levels were assessed by western blotting. DMEM medium without epoximicin was used as a control (**D**) * p<0.05; ** p<0.01, compared to media without epoximicin.

We next examined the kinetics of intracellular BMP2 accumulation following proteasomal blockade. To assess if proteasomal blockade leads to a sustained increase in secreted BMP2 level, time-course experiments were carried out, examining temporal changes in intracellular and secreted BMP2 in response to pharmacological blockade of the proteasomal degradation pathway. Results of this study indicated that addition of epoximicin or MG-132 led to a time-dependent increase in intracellular BMP2 level (**P≤0.01 to *P≤0.05; **Fig 1C** and **Supp. Fig 1B**). Since the duration of transfection can also affect accumulation of the protein from *de novo* synthesis, we also analyzed changes in the intracellular level of BMP2 without epoximicin. Treatment of 293T cells with culture medium, lacking epoximicin did not indicate statistically significant increase in intracellular BMP2 level after 8 hrs of treatment; whereas epoximicin treatment for the same duration led to a significant increase in intracellular BMP2 (**P≤0.01, **Figure 1C**).

Addition of culture medium or medium containing epoximicin medium led to a time-dependent increase in BMP2 secretion into the medium (**Figure 1D**). At each time-point, epoximicin treatment resulted in significantly higher increase in BMP2 secretion in comparison to the increase in BMP2 level from media alone, indicating the role of proteasomal degradation in regulating BMP2 kinetics and secretion (*P≤0.05, **Figure 1D**). Overall, these experiments confirmed that BMP2 expression is regulated by the proteasomal pathway and a pharmacological perturbation of this pathway *in-situ* can be used as a strategy to enhance BMP2 secretion. These experiments further suggest that proteasomal blockade could be an effective strategy to enhance BMP2 level *in situ* and concomitantly enhance its extracellular secretion.

### Proteasomal blockade increases BMP2 half-life through post-translational mechanisms

We next assessed the effect of proteasomal block on steady-state turnover of intracellular BMP2. 293T cells were transfected with a plasmid vector, expressing BMP2 and treated with cycloheximide (Chx) in presence or absence of proteasomal inhibitors. Cycloheximide blocks *de novo* synthesis of proteins, allowing measurement of their steady-state turnover [28]. Proteasomal blockade with either epoximicin or MG-132 led to a significant increase in the intracellular half-life of BMP2 following epoximicin treatment (**Figure 2A**; **P≤0.01 at 30- and 60-min post-treatment with epoximicin and *P≤0.05 at 30 and 60 minutes post-MG-132). Addition of epoximicin and MG-132 led to significant delay in BMP2 degradation, almost doubling BMP2 half-life (**Figure 2A**; 30-min post-cycloheximide addition in comparison to >60 min following addition of epoximicin or MG-132, *P≤0.05 to P≤0.01).

**Figure 2:**
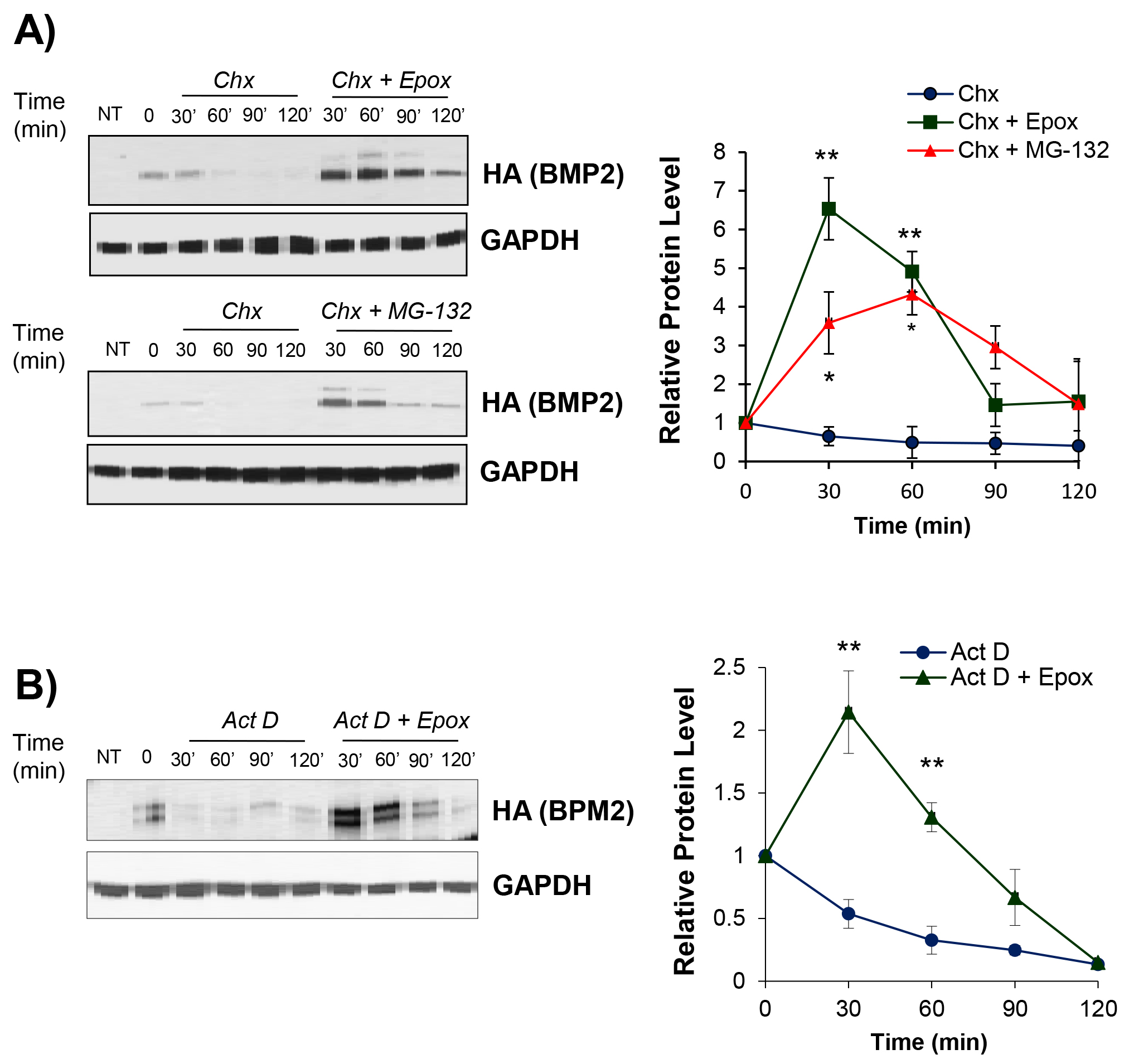
Cycloheximide and actinomycin-D chase experiments to determine the effect of proteasomal blockade on constitutive BMP2 turnover. 293T cells were transfected with p-CMV3-HA-BMP2 and split 24 hrs post-transfection. Twelve-to-sixteen hrs later, cells were treated with 75 μg/mL cycloheximide (*Chx*) alone or along with proteasomal inhibitors; 2 μM epoximicin (*Epox*) or 20 μM MG-132. **(A)** The cells were harvested at indicated times following treatment and changes in BMP2 levels, relative to total cellular protein, were evaluated as a function of time by immunoblotting with anti-HA antibody, followed by densitometric quantitation from triplicate experiments; **p<0.01, *p<0.05, compared to *Chx* treatment alone. (**B**) 293T cells were treated with 10 μg/mL Actinomycin-D (*Act D*) in presence or absence of epoximicin and harvested at indicated intervals following treatment. Cell lysates were analyzed by immunoblotting with anti-HA antibody. Densitometric quantitation from triplicate experiments is shown next to the blot; **p<0.01, *p<0.05, compared to *Act D* treatment alone.

Similarly, BMP2 was rapidly degraded following treatment with Actinomycin-D, an RNA polymerase II inhibitor, which prevents the synthesis of new mRNA transcripts (**Figure 2B**). However, co-treatment with proteasomal inhibitor, epoximicin, significantly delayed BMP2 degradation, suggesting the involvement of post-transcriptional mechanisms in regulating constitutive BMP2 level (**Figure 2B**; **P≤0.01 at 30 and 60 min post epoximicin treatment). Overall, these data suggested that proteasomal blockade increases BMP2 turnover through post-transcriptional and post-translational mechanisms.

### BMP2 is regulated by ubiquitination-mediated proteasomal degradation

Following our finding that BMP2 is regulated through a post-translational mechanism involving the proteasomal pathway, we next investigated what signals BMP2 to this route. Proteasomal degradation is most commonly regulated through ubiquitination-mediated targeting of proteins to the proteasomal pathway. To ascertain if ubiquitination mediates proteasomal targeting of BMP2, 293T cells were co-transfected with BMP2 and HA-tagged ubiquitin expression plasmids and ubiquitination status of BMP2 following a blockade of the proteasomal pathway was analyzed by immunoprecipitation with BMP2 antibody (**Figure 3A**). Following treatment with epoximicin, the cell lysates (input) and the supernatants were also analyzed by western blotting (**Figure 3B**). As shown in Figure 3A, the appearance of distinct HA-smears in BMP2 immunoprecipitates suggested that baseline ubiquitination of BMP2 controls constitutive levels of proteins (**Figure 3A**, lane 4). The HA-smear intensity in immunoprecipitated BMP2 increased following treatment with epoximicin, suggesting an accumulation of polyubiquitinated BMP2 due to blockade in clearance of the ubiquitinated protein through the proteasomal degradation route (**Figure 3A**, lane 5). In agreement with results shown in Figure 1, BMP2 level *in situ* and its secretion into the medium was concomitantly enhanced in BMP2 and ubiquitin plasmid co-transfected cells that were treated with epoximicin, but not in cultures lacking epoximicin (**Figure 3B**). Overall, these experiments further confirmed that intracellular BMP2 turnover is affected by ubiquitination-mediated proteasomal turnover.

**Figure 3:**
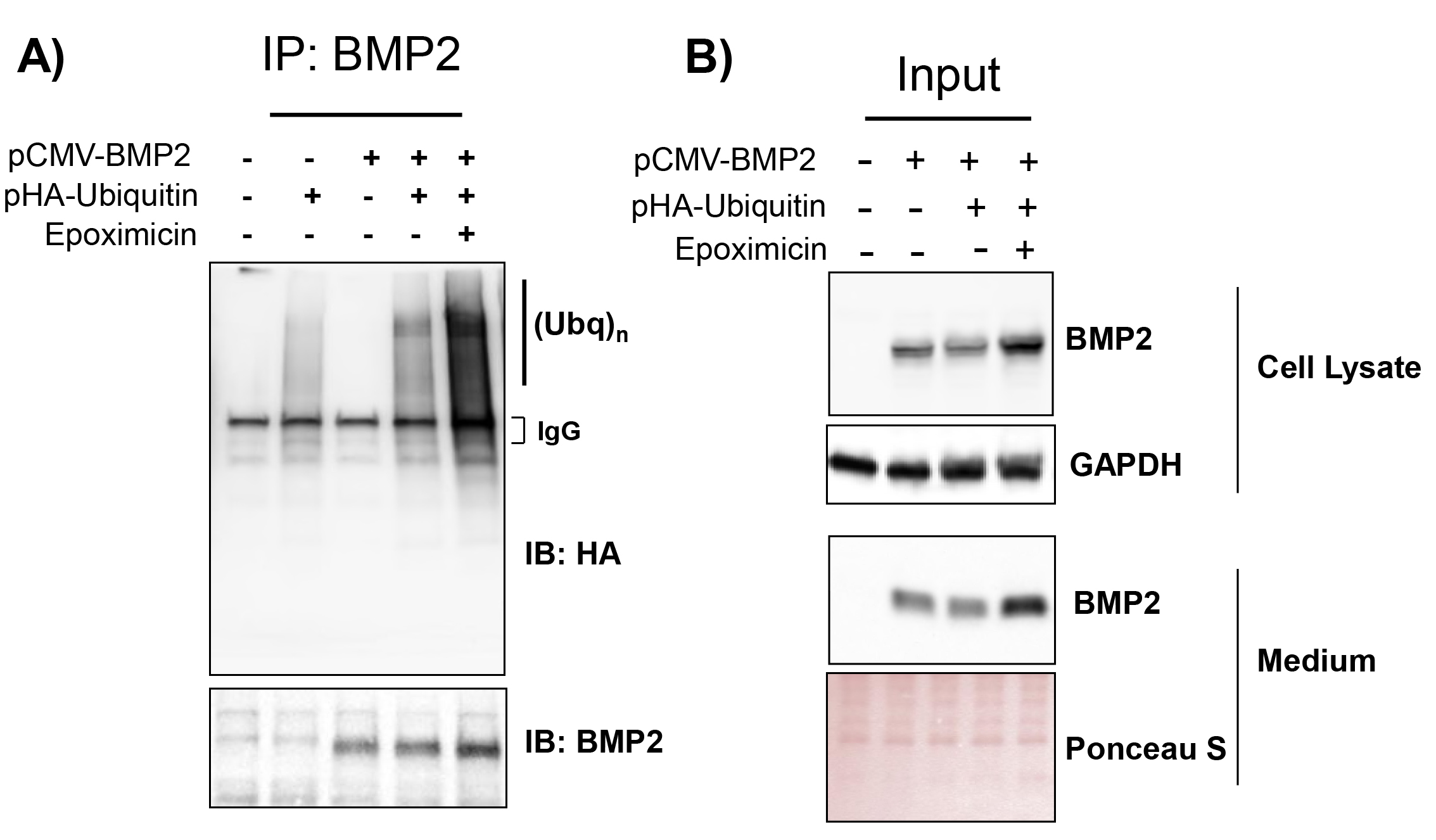
BMP2 undergoes ubiquitination-mediated proteasomal turnover. (**A**) 293T cells were co-transfected with BMP2 and HA-tagged-ubiquitin plasmids. Cells were treated with epoximicin or DMSO vehicle for 1 hr and BMP2 was immunoprecipitated. The immunoprecipitates were separated on 4-20% gradient polyacrylamide gels and changes in BMP2 ubiquitination status were evaluated by immunoblotting with anti-HA antibody. Epoximicin led to a marked increase in HA smear intensity indicating an accumulation of polyubiquitinated BMP2. (**B**) Lysates used for the ubiquitination assay were also analyzed by immunoblotting with BMP2 antibody (upper panel). Media was collected from transfected cells following treatment with epoximicin and assessed by immunoblotting (lower panel).

### Lysine residues at positions 185, 272, 278, and 281 in pro-BMP2 domain and at 290, 293 and 297 within the mature-BMP2 region are critical for BMP2 intracellular turnover and secretion

Since ubiquitination typically occurs on lysine residues, we next sought to identify lysine residues critical for BMP2 proteasomal turnover. Because there were no apparent, pre-defined degradation motifs in BMP2, we conducted site-directed mutagenesis by individually mutating each of the nineteen lysine residue present in BMP2. Lysine-to-arginine mutagenesis is frequently employed to identify sites of ubiquitination by replacing ɛ-amino group on lysine with arginine; wherein such a mutation on potential site renders the protein defective to the attachment of ubiquitin [27, 29]. Of the nineteen lysine residues throughout BMP2, ten are located in the pro-region of the protein and the other nine are located in the mature region of BMP2 that is secreted out of the cell. We first mutated lysine residues within the pro-domain of BMP2, encompassing residues 1 to 281. When compared to wild-type BMP2, lysine-to-arginine substitutions at residues 185, 272, 278, and 281, located within the BMP2 prodomain significantly enhanced constitutive BMP2 expression level and also significantly increased the secretion of the protein into the media (p≤0.05; **Figure 4A,** lower panel). Although point mutations at K64, K127, K178, and K185 increased BMP2 expression intracellularly, a concomitant increase in BMP2 secretion was statistically significant only for mutation on K185. Mutations at K236 and K241 had no influence on either intracellular BMP2 or the level of secreted form.

**Figure 4:**
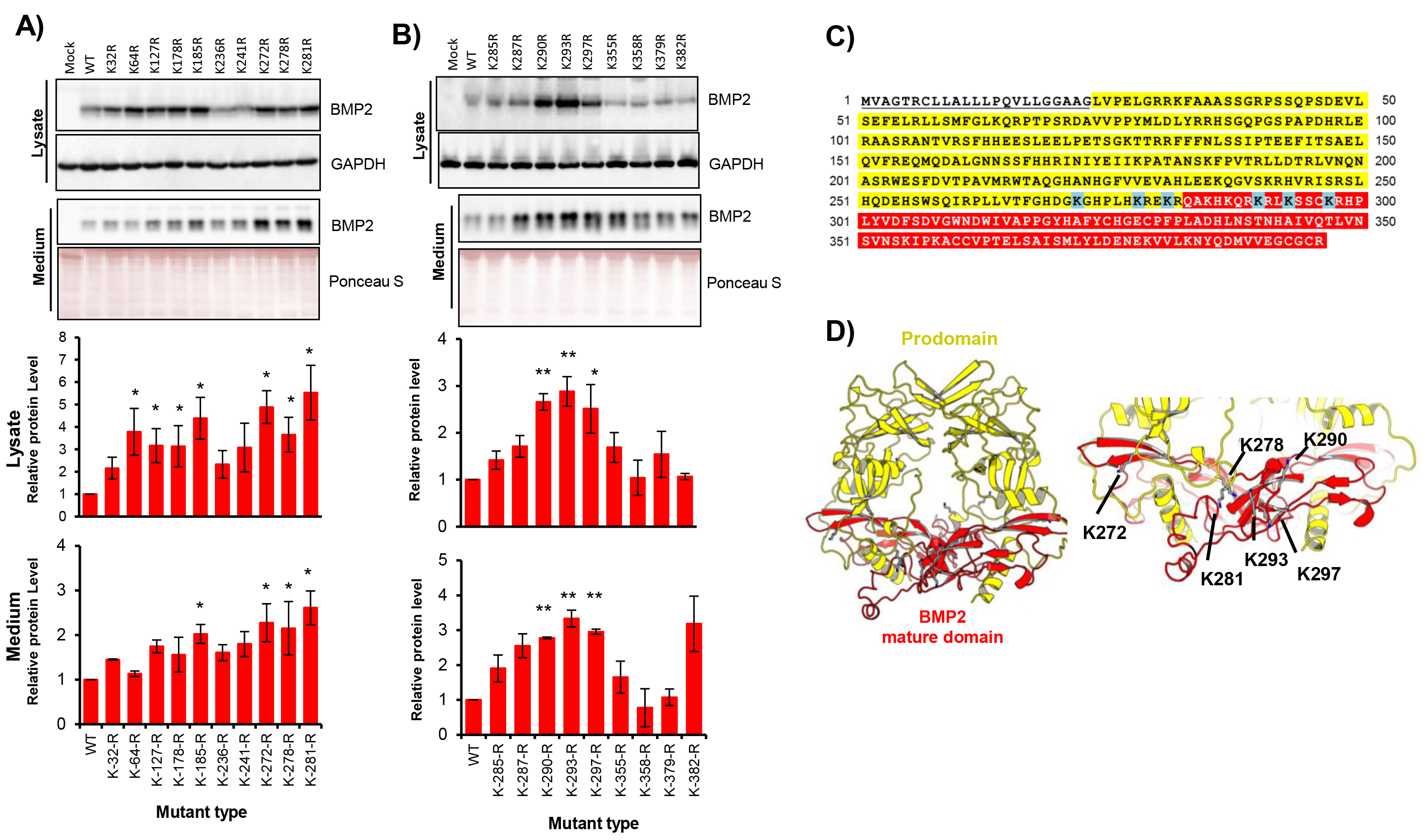
Site-directed mutagenesis on lysine residues present in BMP2. (**A**) Individual BMP2 mutations, substituting lysine residues, spanning the entire protein, were performed by replacing with arginine using an expression plasmid. 293T cells were transfected with plasmid encoding wild-type BMP2 or indicated K-to-R mutations, individually at pro-domain and mature domain. Cell lysates and conditioned were prepared 24 hrs post-transfection and analyzed by immunoblotting to assess changes in BMP2 levels. Analysis of mutations spanning BMP2 prodomain is shown in (**A**) and those spanning the mature BMP2 is shown in (**B**). Densitometry analysis from three independent experiments is provided at the bottom (*P<0.05, **P<0.01 vs. wild-type BMP2). The amino acid sequence of human BMP2 pro-domain is highlighted in yellow, and the sequence of mature BMP2 is highlighted in red. Identified lysine residues, critical for BMP2 intracellular stability are highlighted in blue (**C**). Structural model of BMP2 pro-domain- was generated using a crystal structure of TGF-β1 as a template. Homology model was built using Swiss-Model automated server, and structure figures were generated using PyMol. Pro-BMP2 domain and mature BMP2 domain are highlighted in yellow and red, respectively. Locations of critical lysine residues important for BMP2 turnover are shown in the right (**D**).

We next investigated the lysine residues present in the mature region of BMP2, which is extracellularly secreted and mediates canonical and non-canonical signaling, encompassing residues 282 to 396 (**Figure 4B**). Among 9 lysine residues in the mature-BMP2 domain, mutations at K290, K293, and K297 significantly increased the expression of BMP2 both intracellularly and in its secretion significantly enhanced protein secretion, compared to wild-type BMP2 (**Figure 4B**). Interestingly, homology modeling of pro-domain-associated BMP2 sequence (shown in **Figure 4C**) further indicated that the most influential lysine residues (K272, K278, K281, K290, K293, and K297; highlighted in blue) appear to form a lysine cluster near the cleavage site that releases a mature form of BMP2 (**Figure 4D** and **Supp. Figure 2**).

### Lysine-to-arginine substitutions that prevent *in-situ* proteasomal degradation do not affect the bioactivity of mutant BMP2

To confirm that the K-to-R mutations do not compromise the biological activity of BMP2, we next assessed the ability of mutant BMP2 to induce canonical Smad phosphorylation, as compared to wild-type BMP2 (**Figure 5**). Conditioned media was collected from 293T cells, transfected with either WT or mutant BMP2 plasmids individually, and was used to test Smad phosphorylation in MG-63 cells through BMP receptors. Results of this experiment indicated potent and comparable Smad phosphorylation in MG-63 cells from lysine-to-arginine mutants at K185, K272, K281, K290, and K293, suggesting mutations on identified lysine residues do not compromise the functionality of the molecule (**Figures 5A & B**).

**Figure 5:**
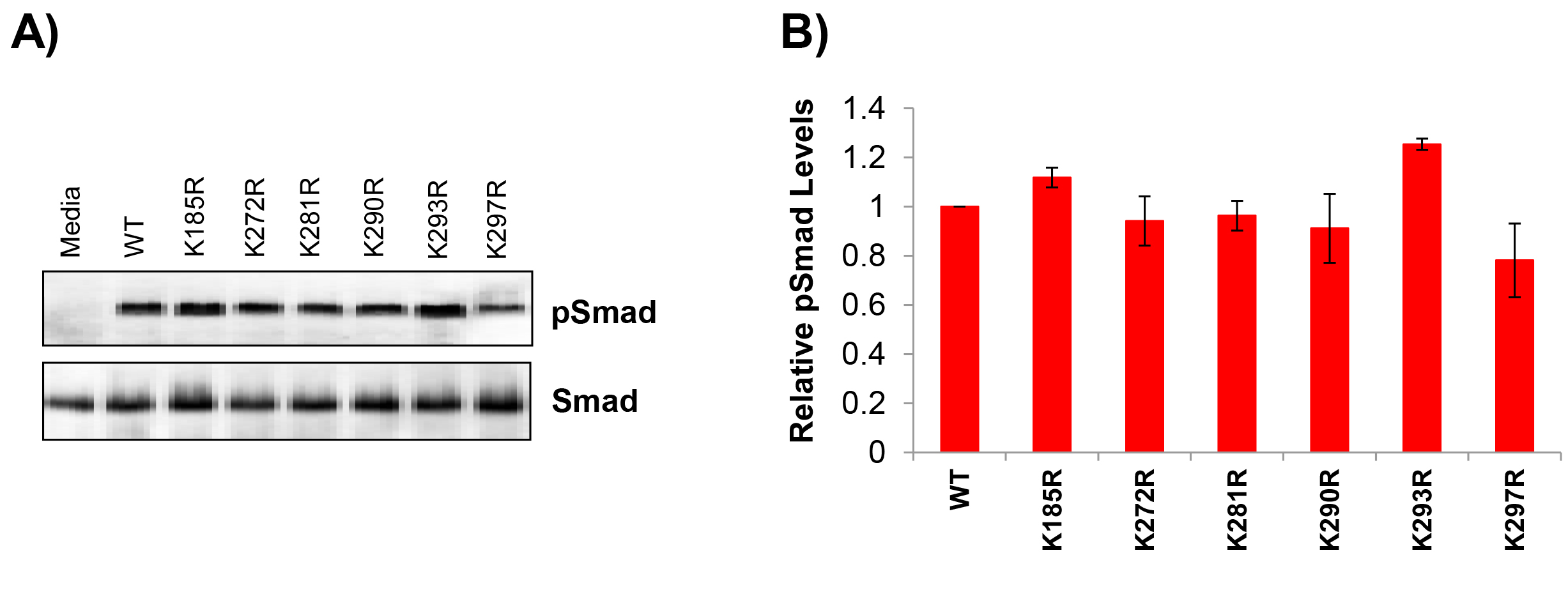
Smad phosphorylation assay to determine the biological activity of BMP2 with lysine-to-arginine substitution mutants. (**A**) 293T cells were transfected with plasmids encoding wild-type or indicated lysine-to-arginine mutant BMP2. Twenty-four hrs post-transfection, cultures were replenished with serum-free medium and cells were incubated for 16 hrs prior to harvesting conditioned media. Media were tested on MG-63 cells for activation of BMP2 canonical signaling pathway. MG-63 cells were serum-starved overnight prior to treatment with conditioned medium from BMP2 plasmid transfected 293T cells. After 1 hr of treatment, MG-63 cells were harvested, lysates prepared and analyzed by western blot using phospho-Smad and total Smad antibodies. (**B**) Densitometric quantitation of respective P-Smad levels, relative to total Smad levels, from triplicate experiments is shown.

## DISCUSSION

The present study sought to identify putative mechanisms regulating constitutive turnover of intracellular BMP2 and how they influence its extracellular secretion. Although transcriptional regulation and proteolytic processing of BMP2 has been investigated in detail, studies dissecting its intracellular degradation and its effect on extracellular secretion are lacking [14, 30-33]. Half-life of BMP2 in circulation is very short, averaging 7-16 min systemically [11]. Many studies have also documented a significantly increased risk of clinical complications due to supraphysiological amounts of protein in currently used BMP2 products [4, 6-10]. Therefore, there is a pressing need to identify mechanisms that modulate BMP2 turnover to improve BMP2 delivery methods without adverse effects [6, 7, 10, 13, 34].

We hypothesized that minimizing BMP2 degradation *in-situ* would improve its intracellular retention and enhance extracellular secretion and bioavailability [35]. Since the constitutive turnover of a majority of cellular proteins is largely governed by proteasomal degradation, we first examined the effect of proteasomal blockade on BMP2 turnover. Pharmacological blockade of the proteasomal pathway by epoximicin and MG-32 led to a significant increase in the intracellular retention of BMP2 and more importantly, led to a concomitant enhancement in its secretion. Since the increase in BMP2 could have been due to an increase in BMP2 synthesis through alternative mechanisms that are also regulated by the proteasomal pathway, we performed cycloheximide chase experiment to determine if BMP2 is regulated at the level of synthesis or post-translationally by proteasomal degradation [14, 27, 36-38]. Results of this study confirmed that BMP2 levels were stabilized with proteasomal inhibitors even after transcription and translation steps were blocked, suggesting the existence of direct post-translational mechanisms mediating the increase in BMP2 level [36].

The proteasomal pathway is regulated through a wide variety of post-translational modifications, ubiquitination being the most frequent signal for shunting proteins to the degradation pathway [15, 39]. BMP2 is also known to be post-translationally modified and there is substantial evidence that other members of BMP/TGF family undergo post-translational modifications [17-22]. Glycosylation of BMP1 and BMP2, for instance, is known to regulate turnover and secretion of these ligands [17, 18]. BMP2 is known to be glycosylated on N135, mutation of which results in retention of the protein within the endoplasmic reticulum [17]. High-throughput proteomic studies have suggested that other members of this family such as TGFβ1, BMP3, BMP4, BMP6, BMP7, and BMP8 are indeed ubiquitinated [20-22]. However, there have been no systemic studies to ascertain whether BMP2 is ubiquitinated and if ubiquitination of BMP2 or plays a role in its cellular turnover. To address this, we carried out ubiquitination assay and identified that under physiological conditions, BMP2 is regulated by ubiquitination-mediated turnover. Garrett *et al* demonstrated that proteasomal inhibitors stimulated BMP2 mRNA expression by inhibiting the proteolytic processing of Gli3 protein, a transcriptional factor necessary for BMP2 synthesis [23]. However, evidence of BMP2 ubiquitination and direct regulation of the protein by the proteasomal pathway was not suggested in this study [23]. In light of these studies, it is important to consider that the cellular abundance of BMP2 maybe regulated through its direct degradation via the proteasomal pathway as well as through other BMP2-modulating proteins that are regulated by this degradation route.

To overcome problems relating sub-optimal BMP2 delivery due to rapid clearance of the protein, many groups including ours have employed strategies to optimize the dose of BMP2 at the site of bone defects [13, 14, 34]. For instance, polyglutamate (E7) peptide-directed coupling of BMP2 to various scaffolds and the use of shortened BMP2 knuckle epitope peptide has shown promise in optimizing BMP2 dose at the site of bone defects [40, 41]. Although these studies are promising, issues related to intracellular turnover of BMP2 still remain critical. Hence, we sought to address this important question by identifying which site(s) within the protein is responsible for its intracellular turnover. Lysine-to-arginine mutagenesis is frequently employed to identify sites of ubiquitination by replacing ɛ-amino group on lysine with arginine; wherein such a mutation on potential site renders the protein defective to the attachment of ubiquitin [29]. We reasoned such a conservative substitution would reduce the susceptibility to BMP2 to ubiquitination and reduce its intracellular loss, in turn, increasing its availability for secretion. We found that mutating C-terminal lysine residues at positions 272, 278, and 281 within the pro-domain and residues 290, 293, and 297 in the mature BMP2 significantly increased intracellular BMP2 and its subsequent secretion, without compromising the biological activity of the protein. It is also noteworthy that our structural modeling of prodomain-associated BMP2 indicated that these ubiquitination-susceptible K272, K278, K281, K290, K293, and K297 residues are clustered near the proprotein convertase S1 cleavage site (REKR) that releases a mature form of BMP2, suggesting the possibility that proteasomal regulation of both the full-length protein (residues 1-396) as well as proteolytically-processed mature BMP2 occurs predominantly inside the cell. Further study is warranted to identify if such lysine clusters in other members of TGF beta superfamily have a role in proteasomal regulation of ligands.

Vallejo *et al* has previously shown that BMP2 folding and dimerization is a very slow process involving at least one long-lived intermediate [42]. Other studies have indicated that co-translational degradation of partially folded protein intermediates may occur during degradative mechanisms during protein quality control [43]. It is conceivable that a fraction of BMP2 is ubiquitinated during its post-translational processing, folding and proteolytic processing; allowing cognate ubiquitin ligases to access residues in a partially-folded intermediate, which may otherwise be sequestered in the interior of the protein. It should be noted that the cellular degradation profile of BMP2 is independent of its systemic degradation. In this regard, it is important to consider that blocking proteasomal degradation of BMP2 would increase its intracellular abundance without affecting its systemic status. However, in gene, or genetically-engineered cell therapy methods, an increased cellular turnover and a concomitant increase in secreted BMP2 level would greatly benefit in bioavailability of this osteogenic factor, overcoming issues related to sub-optimal delivery and heterotopic ossification. The lysine mutants of BMP2 that we identified and validated retain signaling functions comparable to wild-type protein offers a new approach to overcoming existing limitations in using this osteogenic factor. It remains to be tested if mutating all of the identified key lysine residues or those within either the prodomain or in the mature protein would render further benefit to this approach. Overall, our study uncovers previously unreported mechanisms of BMP2 regulation, which could be potentially used in clinical applications.

## MATERIALS AND METHODS

### Cell lines and Reagents

Human embryonic kidney cell line, 293T, and human osteosarcoma cell line, MG-63 were obtained from American Type Culture Collection (Manassas, VA) and maintained in DMEM media (Life Technologies; Grand Island, NY, Cat # 11965-092) containing 10% fetal bovine serum (Omega Scientific; Tarzana, CA, Cat # FB-02) and 100 units/ml penicillin-streptomycin antibiotic (Life Technologies; Grand Island, NY, Cat # 15140122). MG-132 and Epoximicin were obtained from ApexBio (Houston, TX, Cat # A2585 and A2606 respectively). Cycloheximide, and Actinomycin D was purchased from Sigma-Aldrich (St. Louis, MO, Cat # C7698) and Tocris Biosciences (Minneapolis, MN, Cat # 1229), respectively.

### Transfections

The HA-tagged BMP2 expression plasmid, pCMV3-HA-BMP2, was obtained from Sino-Biological (Beijing, China, Cat # HG10426-NY). This CMV promoter-driven mammalian-expression plasmid encodes for HA epitope-tagged BMP2 (NCBI Ref Seq NM_001200.2, 1191 bp) and allows for transient and stable expression of full-length BMP2 and concomitant secretion of mature BMP2 into the extracellular medium. 293T cells were transfected either with calcium phosphate or with PolyJet^™^ *in vitro* DNA transfection reagent (SignaGen Laboratories: Rockville, MD, Cat # SL100688) according to manufacturer’s instructions [44]. The cells were harvested 24-36 hrs following transfection for downstream protein assays as described below. Protein expression of the full-length protein in the cell lysate, and the mature BMP2 in the cell culture media was confirmed by immunoblotting.

### Cycloheximide and Actinomycin D chase assays

To compare the turnover of BMP2, pCMV3-HA-BMP2 plasmid was transfected into 293T cells as described above. Following transfection, cells were pre-treated with proteasomal inhibitor epoximicin (2 μM) or MG-132 (20 μM) for 60 min. After treatment, cells were briefly washed with PBS and exposed to cycloheximide (75 μg/mL) in presence or absence of proteasomal inhibitors. Cells were then harvested at regular intervals and lysates from each condition were analyzed by immunoblotting with anti-HA or anti-BMP2 antibody to measure changes in post-translational turnover following proteasomal block. Actinomycin D (10μg/mL) was used when assessing post-transcriptional changes.

### Ubiquitination assay

pRK5-HA-ubiquitin plasmid (Addgene; Cambridge, MA, Cat # 17608) was co-transfected with BMP2 plasmid into 293T cells. Cells were then treated with proteasomal inhibitors or with DMSO for 2 hrs and then harvested. Lysates from harvested cells, containing 500-1000 μg protein, were subjected to immunoprecipitation (IP) with 1-2 μg of BMP2 antibody using the ImmunocruzCruz^TM^ F kit (Santa Cruz; Dallas, TX, Cat # 45043) according to the manufacturer’s instructions. The precipitates were separated on 4-20% polyacrylamide gradient gels (Bio-Rad; Hercules, CA, Cat # 4561093) then transferred to nitrocellulose membranes (Bio-Rad; Hercules, CA, Cat # 162-0115) by overnight wet transfer and subjected to immunoblotting with HA-antibody as described below to assess changes in ubiquitinated BMP2.

### Site-directed mutagenesis

The Quik-Change^TM^ site-directed mutagenesis kit (from Agilent Technologies (Santa Clara, CA, Cat # 210518) was used to generate all BMP2 mutants. Briefly, DNA primers (see **Supplementary Table 1**) were employed to introduce single nucleotide changes (A**<u>A</u>**A to A**<u>G</u>**A; A**<u>A</u>**G to A**<u>G</u>**G) within the lysine codons present in the BMP2 coding region. All constructs were sequenced and verified prior to assay.

### Homology modeling of prodomain-associated BMP2

To illustrate potential ubiquitination sites, the structural model of full-length BMP2 (residues 24-396, See also **Figure 4C** and **Supp. Figure 2**) was generated using SWISS-Model server [45]. Among top-scoring searched models, the crystal structure of prodomain-associated TGF-beta1 (PDB: 3RJR) was used as a template for homology model building. All the figures were generated using the PyMol program (Schrödinger, Cambridge, MA, USA; http://www.pymol.org).

### Western blot analysis

Cells were harvested following each transfection and lysed with 1X RIPA buffer, supplemented with protease and phosphate inhibitors (Thermo Fisher Scientific, Waltham, MA, Cat # 89900, A3278428, and A32953, respectively). Lysates (25-40 μg) were separated on Mini-PROTEAN^TM^ (Bio-Rad, Hercules, CA) and transferred on to nitrocellulose membranes. The membranes were blocked for 1 hour with 5% milk in 0.05% TBST and probed with the following antibodies at 4°C overnight: anti-BMP2 monoclonal detection antibody conjugated to horseradish-peroxidase (Sino-Biological, Beijing, China, Cat No SEK 10426, dilution - 1:5000 - 1:10000), anti-HA antibody (Thermo Fisher Scientific, Cat # 14-6756-63, dilution 1:0000), Phospho-Smad1/5/9 (D5B10) Rabbit mAb (Cell Signaling, Danvers, CA, Cat No 13820T, dilution 1:500), Smad1 (D59D7) XP^®^ Rabbit mAb (Cell Signaling, Danvers, CA, Cat No 13820T, dilution 1:1000). Blots were developed using the corresponding horseradish peroxidase-conjugated secondary antibodies with Immobilon^TM^ chemiluminescence HRP substrate (Millipore, Billerica, MA, Cat # WBKLS0500) and the images were obtained with a PXi gel imaging system (Syngene, Frederick, MD, USA). Densitometric analysis was carried out using NIH-ImageJ^TM^. Protein expression levels of all experimental conditions were represented as relative values (means ± SEM) of controls.

### Smad phosphorylation assay for functional validation of BMP2 variants

To assess BMP2 bioactivity, downstream Smad phosphorylation was performed in MG-63 osteosarcoma cell line, as previously described [46]. Briefly, MG-63 cells were treated with serum-free conditioned medium from 293T cells transfected with mammalian expression vectors encoding either the wild-type BMP2 or those containing lysine mutations. Prior to addition of conditioned media, MG-63 cells were serum-starved overnight. Cells were harvested one hr post-treatment and Smad-phosphorylation was evaluated in cell lysates by immunoblotting with Phospho-Smad1 (Ser463/465)/ Smad5 (Ser463/465) rabbit monoclonal antibody (clone D5B10; Cell Signaling, Danvers, CA, dilution 1:500). Smad-phosphorylation was normalized with total Smad levels, using total Smad antibody (Cell Signaling, Danvers, CA, Cat No 13820T, dilution 1:500).

### Statistical analysis

The statistical significance of data, obtained from the quantitative analysis of protein expression, was determined by Student’s t-test. Values with P<0.05 were considered significant.

## ACKNOWLEDGEMENTS

This work was supported by U.S. National Institute of Health/ National Cancer Institute Grants R01AR060948, R01CA184770 (S.P.), and Department of Defense Grant PR141945.

## AUTHORSHIP CONTRIBUTIONS

VK, JHL, HW, and SB performed experiments, VK, and JHL analyzed data, VK, J.H.L wrote the manuscript; S.P. designed work, provided resources, analyzed data and wrote manuscript.

